# Reconstructing contact network structure and cross-immunity patterns from multiple infection histories

**DOI:** 10.1101/701946

**Authors:** Selinger Christian, Alizon Samuel

**Affiliations:** MIVEGEC, Univ. Montpellier, CNRS, IRD, Montpellier, France.

## Abstract

Interactions within a population shape the spread of infectious diseases but contact patterns between individuals are difficult to access. We hypothesised that key properties of these patterns can be inferred from multiple infection data in longitudinal follow-ups. We developed a simulator for epidemics with multiple infections on networks and analysed the resulting individual infection time series by introducing the concept of infection barcodes. We find that, depending on infection multiplicity and network sampling, infection barcode summary statistics can recover network properties such as degree distribution. Furthermore, we show that by mining infection barcodes for multiple infection patterns, one can detect immunological interference between pathogens (i.e. the fact that past infections in a host condition future probability of infection). The combination of individual-based simulations and barcode analysis of infection histories opens promising perspectives to infer and validate transmission networks and immunological interference for infectious diseases from longitudinal cohort data.

**Author summary:** Infectious disease dynamics are constrained both by between-host contacts and pathogen interactions within a host. Furthermore, multiple parasites circulate such that hosts are infected (sometimes simultaneously) by a variety of strains or species. We hypothesise that multiple infection history can inform us about the networks on which parasites are transmitted, but also on within-host interactions such as immunological interference. We developed a simulator for multiple infections on networks. By combining intuitive novel metrics for multiple infection events and established tools from computational data analysis, we show that similarity in infection history between two hosts correlates with their proximity in the contact network. By analysing pathogens co-occurrence patterns within hosts, we also recover immunological interference at the population level. The demonstrated robustness of our results in terms of observability, network clustering, and pathogen diversity opens new perspectives to extract host contact and between-pathogen immunity information from longitudinal cohort data.

## Introduction

Host populations are often assumed to be ‘well-mixed’ even though individuals tend to only interact with a small subset of the whole population and these contact patterns between individuals can dramatically affect the way epidemics spread [1–3]. For instance, it was shown during the early phase of the HIV pandemics that the average number of sexual partners but also the variance in the number of partners both increase the basic reproduction number (*R*_0_) of sexually transmitted infections [4]. Since then, studies have identified how key parameters of the host contact network affect the risk of outbreak [5, 6] and the spread of an epidemic [7–13].

Network reconstruction, *i.e.* the inference of adjacency weights based on observations of a dynamical system acting on the network, is a well-established research topic in engineering [14], and has recently been applied to infectious disease dynamics [15–17]. In the field of epidemiology, measuring contact networks is a lively research topic [18], ranging from definition issues (defining a ‘contact’), to assessing the appropriateness of various types of data. For livestock [19], interactions between interconnected farms can be well approximated through shipping logs. Other settings, such as wild populations or human populations, are more challenging to analyse. For humans, the field has traditionally relied on self-reported data, but new insights have been provided by airline transportation registries [20] or cell phone data [21]. More recently, parasite sequence data was used to infer network properties by analysing phylogenies of infections [22]. Importantly, this genetic data, which by definition pre-dates new outbreaks, can be used to make relevant predictions [23]. Note that we use the terms parasite and pathogen interchangeably encompassing both micro- and macro-parasites.

We hypothesise that host contact network properties can be inferred from individual longitudinal data about infection status by taking advantage of multiple infections as a unique descriptor of a host’s position within the network. Such longitudinal data is classically used in epidemiological studies to measure the odds that a specific event may occur [24], however, they are rarely coupled to mathematical models of disease spread.

To track multiple parasite strains or species in these individual histories is both feasible and highly relevant, as the increasing affordability and power of sequencing strengthen the assertion that most infections are genetically diverse [25]. In fact, in the case of genital infections by human papillomaviruses (HPVs), not only do we know that coinfections, i.e. the simultaneous infection by multiple genotypes, are the rule rather than the exception, we also know that they strongly correlate with the number of lifetime sexual partners [26]. Another classical example is the distribution of the number of macroparasites per host, which has been used to infer population structure [27]. This makes multiple infections an ideal candidate to measure epidemic properties but also detect potential within-host interactions between parasite strains or species [28, 29].

To show how individual infection histories for multiple parasite strains can inform us about the underlying transmission contact network, we conduct a simulation study, which requires alleviating two obstacles. First, we need to simulate multiple infections on a network, a task few studies have attempted [12, 30–35]. For this, we take advantage of recent developments in stochastic epidemiological modelling and implement a non-Markovian version of the well-known Gillespie algorithm [36]. This allows us to make sure a host’s infection history is not lost every time it acquires a new strain. It also addresses statistical evidence that for many parasites infection duration does not follow exponential but heavy-tailed distributions [37–42]. The second main obstacle resides in extracting information from longitudinal follow-up data. To compare these infection histories, we use barcode theory from computational topology [43].

We integrate infection histories into our multiple infection modelling framework by accounting for the documented fact that recovery from infection becomes more likely with increasing ‘age’ of infection [44–47]. Then, we simulate individual multi-strain infection histories for epidemiological models with genotype-specific immunity on random clustered networks to demonstrate the impact of network topology on infection diversity. We refer to ‘genotypes’ to describe the parasite diversity, but our results apply to different genotypes of the same species or of different species of parasites provided that their mode of spreading (e.g. airborne, sexually transmitted) on the host network is the same. We include immunological interference between genotypes in our model by constraining the probability of acquiring a new infection in terms of the host’s infection history, akin to hemagglutination inhibition assays (e.g. multi-season influenza strains [48]).

Finally, we show that infection barcodes can inform us both about the connectivity in the network and immunological interference between genotypes in several ways. First, similarity matrices between individual infection histories highly correlate with the network adjacency matrices. Second, an individual host’s network connectivity can be inferred from its connectivity in terms of infection barcodes. Third, by quantifying genotype co-occurrence based on infection histories, we recover model inputs for immunological interference. Fourth, mining frequent infection sequence motifs allows quantifying patterns induced by immunity settings in the spirit of more widely used cross-sectional prevalence surveys.

Taken together, in our simulation study we provide proof-of-concept to reconstruct network and immunity characteristics from novel summary statistics based on multiple infection histories. The demonstrated robustness and limitation of our approach towards network properties, host population sampling and genotyping opens novel avenues towards computational epidemiology of multiple infections.

## Materials and methods

### Simulation algorithm

We developed an event-driven stochastic model of multiple infections on networks in Python. For the purpose of this simulation study, we considered static, random networks to model contact (i.e. edges with binary weights) between hosts (i.e. nodes). Contact networks were generated using the class of random clustered graphs [49–51] implemented in the networkx package in Python [52]. Given a propensity list of edge degrees and triangle degrees as input, this algorithm generates a random graph with predefined average degree and average clustering coefficient. The clustering coefficient of a graph was defined as the local clustering coefficient (i.e. the degree of a node divided by the number of all possible edges in the node’s neighbourhood) averaged over all nodes. Degree dispersion was defined as the ratio of degree variance and degree mean. Degree assortativity (i.e. the propensity for nodes in the network that have many connections to be connected to other nodes with many connections) was defined following [53].

Upon network initialisation, we randomly seeded outbreaks of one infection per genotype, allowing for up to four genotypes per host. Disease dynamics followed Gillespie’s stochastic simulation algorithm (SSA) [54, 55] adapted to a non-Markovian setting [56] to incorporate memory-dependent processes. Here, in particular, we considered recovery from infection as a process depending on the age of infection.

The simulations also feature potential immunological interference between genotypes. This was defined by an immunity input matrix *I_ij_* ∈ [0, 1], where *I_ij_* is the probability not to acquire an infection with genotype *i* given prior infection with genotype *j*, i.e. upon exposure from an infectious edge with genotype *j*. We sampled independent Bernoulli random variables with probabilities *I_ij_* , for all genotypes *i* contained in the infection history of the exposed host. In these simulations, we explored five immunity settings defined as follows: 

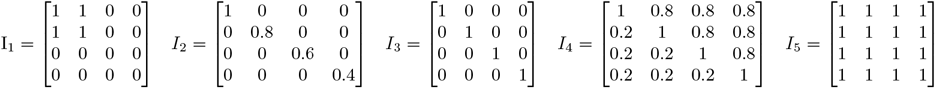

The matrix *I*_1_ corresponds to partial cross-immunity, with sterilising immunity between the first two genotypes, *I*_2_ models decreasing homologous immunity, where genotype *g*_1_ induces sterilizing immunity, whereas immunity is decreasing for the remaining genotypes. *I*_3_ models homologous immunity, i.e. when each genotype induces sterilizing immunity to future infection with the same genotype. *I*_4_ is an example for asymmetric cross-immunity, i.e. immunity to infection with *g*_1_ after clearing an infection with *g*_2_ is stronger than immunity to infection with *g*_2_ after clearing an infection with *g*_1_. Finally, *I*_5_ models sterilizing cross-immunity. Transmission and recovery rates were considered equal for all genotypes and fixed in advance.

The Gillespie SSA allowed us to simulate disease dynamics by performing regular updates in the values of the rates. For this, at each time point, we first created a rate vector {*r_k_}_k∈E_* indexed by the set of all possible events *E* (recovery from an ongoing infection or acquisition of a new infection from an infectious host in the graph neighbourhood). If the node *i* had spent *t_i,g_* time in an infection with genotype *g*, then the rate of recovery from this infection was assumed to be given by a Weibull hazard rate function 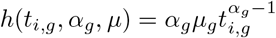 with shape parameter *α_g_* and scale parameter *μ_g_*. The default setting for recovery was *α_g_ =* 2 and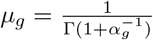 such that the mean was equal to one, while the probability of clearance increased with the age of infection, motivated by infection duration literature (see Introduction). For normalisation purposes, we assumed that the rate of infection for any node currently not infected with genotype *g* was unity, such that the average number of secondary infections was uniquely determined by the average node degree.

To better handle the computational workload of multiple infections on networks, we stored the infection histories in an interval tree data structure containing the times of onset and end of each infection episode, the genotype, and the host number (this is an individual-based model and not an event-based model). We refer to this data structure as infection barcodes (see below). At a given time, from the rates vector, we first drew an exponentially distributed random variate with rate *k r_k_* to determine the time increment to the next event. Certain rates *r_k_* depended on the infection age but, as shown in [56], drawing exponentially distributed random variables with respect to these rates provides good Markovian approximations of non-Markovian processes, granted the number of events is sufficiently large. We then drew a random variate according to the total probability vector {*r_k_*/Σ_*k*_ *r*_*k*_} _*k∈E*_ to determine the nature of the next event. Depending on the event, we updated the list of possible events. In the case of recovery of host *i* from infection with genotype *g*, we wrote the end time of the infection episode to the interval tree, removed the recovered infection from the rate vector, and added potential infections with genotypes other than *g* from the host’s network neighbourhood. In the case where a host *i* is newly-infected with *g*, we wrote the time of onset of the infection episode to the interval tree, removed all possible edges from *g*-infected neighbours of *i*, and added possible infection edges from *i* to all neighbours that had not been or were not currently infected with *g*. We then updated the rates vector again and proceeded until the epidemic became extinct.

Unless stated otherwise, we simulated disease transmission with four genotypes introduced simultaneously on the giant component of random clustered graphs with 250 nodes with average degree of 4 and an average clustering coefficients of 0.34 (referred to as ‘clustered’ networks). For a given parameter set, we seeded a random outbreak at the beginning of each simulation with one infection per genotype and ran 50 stochastic replicates until the disease-free state or a pre-defined time horizon was reached. Epidemic outcomes were reported as average and 95% confidence intervals for equidistant time bins.

The source code and configuration files used for simulations will be publicly available on https://gitlab.in2p3.fr/ETE/MultiNet.

### Genotype diversity and multiplicity of infection

We measured genotype diversity during the course of epidemics with *N =* 4 genotypes by Shannon’s diversity index [57] defined as *H*(*t*) = exp (*i p_i_*(*t*) log *p_i_*(*t*)), where *p_i_*(*t*) is the frequency of infections with genotype *i* relative to all infections present at time *t*. The index *H* ∈ [1, 4] is maximised when all genotypes have equal frequency and minimised when only one genotype is present. We emphasise that this index pertains to diversity at the population-level and does not distinguish between a multiple infection within a single host and single infections in several hosts. To highlight the importance of multiple infections, we also considered the multiplicity of infection (MOI) given by the number of genotypes present within infected hosts at a given time. We reported MOI averaged over all infected nodes.

### Infection barcodes and network properties

We summarised each individual multiple infection history by a barcode, i.e. the set of intervals describing the onset and clearance of infection episodes accumulated within a host during the course of an epidemic. The notion of barcode first arose in computational topology [58] to succinctly summarise and compare topological properties of metric spaces. Mathematically, the infection barcode of a host *A* is a set of triples *A =* {*A*_1_,… , *A_n_*}, where each *A_i_ =* (*b_i_, d_i_, g_i_*) defines the birth 0 ≤ *b_i_ <* ∞ (i.e. infection onset) and death 0 ≤ *d_i_ <* ∞ (i.e. infection clearance) with a particular genotype *g_i_*. We explicitly included the presence of multiple genotypes in a host’s infection history, i.e. a host can simultaneously have several infections, also the same genotype can appear several times in the course of an epidemic. To compare infection histories between hosts, we considered the metric space of infection barcodes endowed with two complementary notions of distance.

In the first approach, we considered a similarity index [59] such that two hosts with largely overlapping infection episodes tended to be similar to each other, regardless of the respective genotypes. More precisely, a given pair of barcodes between any two hosts *A* = {*A_i_*} and *B* = *B_j_* was transformed into a weighted bipartite graph. The nodes of this graph consisted of infection episodes and these nodes were partitioned according to the two hosts. The edge weight of the graph was given by the overlap length 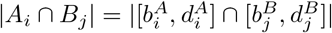 between the infection episodes constituting the edge (*i, j*), taking also into account overlaps between distinct genotypes. To obtain the similarity index *s*_1_(*A, B*), we calculated the maximum bipartite graph matching of this graph. The similarity index was transformed into a distance by *d*_1_ = −2*s*_1_ + Σ_*i*_|*A_i_*| + Σ_*j*_|*B_j_*|.

For the second approach, we considered the overlap length of infection episodes between two hosts with matching genotype *g*. By summing over all genotypes we obtained the similarity index. This 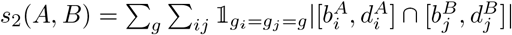 index was transformed into a distance by setting *d*_2_ = −2*s*_2_ + Σ_*i*_ |*A_i_*| + Σ_*j*_ |*B_j_* |.

To determine whether similarity between infection histories implies proximity in the transmission network, we compared the metric space of infection barcodes to the metric space of the network, restricted to nodes that had been infected during the course of the epidemic. For the network, we considered two different distance notions, i.e. the discrete metric based on the binary adjacency matrix and the shortest path distance of the graph.

We used two-sided p-values from the Mantel test [60] between the barcode similarity (resp. distance) matrix and the adjacency or shortest path length matrix respectively to assess for each model the correlation between infection barcode and network topology [61]. Since Mantel permutation tests have been scrutinised for underestimating type I errors in the presence of spatial autocorrelation [62–64], we tested adjacency and shortest path matrices for auto-correlation, using the R package statGraph [65]. While adjacency matrices were not significantly auto-correlated for most lags, the shortest path matrices had significant auto-correlations at all lags (see S5 Fig).

To assess whether spatial correlations between similarity matrices also translated to local properties such as a host’s node degree (i.e. number of network neighbours) or its localisation within the network (i.e. the sum of shortest path lengths to other hosts), we developed a measure of infection barcode connectivity and performed regression analyses of network characteristics on infection connectivity. More precisely, for a given simulation with maximum length *L*, *N_g_* different genotypes and *D +* 1 nodes in the giant component, the normalized infection barcode connectivity of a host *n* was defined by

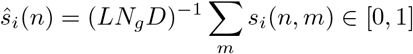

for *i =* 1, 2, i.e. the sum of all barcode similarity values within the network of infected hosts relative to the maximum possible barcode similarity. Since shortest path length is a continuous measure, we performed linear regression using infection barcode connectivity as a regressor and assessed significance for coefficients by p-values from two-sided Z-tests for each of the immunity and network clustering settings. The relationship between node degree and infection connectivity was assessed using a multinomial logistic regression model defined by

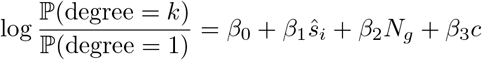

where *β*_0_ is the intercept, *β*_1_ is the coefficient for continuous variable of the normalised barcode similarity degree, *β*_2_ is the coefficient for the categorical variable of *N_g_ =* 2 relative to the base level *N_g_ =* 4, and *β*_3_ is the coefficient for the categorical variable of clustering coefficient *c =* 0.17 relative to the base level *c =* 0.34. We evaluated the model for genotype-agnostic *ŝ*_1_ and genotype-specific *ŝ*_2_ similarity degrees for each of the immunity settings. The multinomial model was trained on 80% of the stochastic replicates and test against the remaining 20%, the area under the curve (AUC) from multinomial precision call curves [66] was used to evaluate predictive power with the R package pROC.

As an alternative, we also explored ordered logistic models defined by

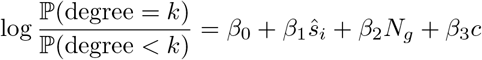

which yielded similar predicted values at consistently higher AIC than multinomial models (data not shown).

### From infection barcodes to immunity

In order to determine immunological interference between genotypes based on individual infection histories, we quantified non-overlapping co-occurrence *C*(*G, H*) of any two genotypes *G* and *H* within the host population. Large *C* values indicate that the presence of genotype *G* in the infection history did not preclude subsequent infection with *H*, i.e. the immunological interference between the two genotypes was weak. Conversely, if *C* is zero, then infection with *G* did prevent infection with *H*, such that *G* conferred full cross-immunity to *H*. This can be written mathematically as

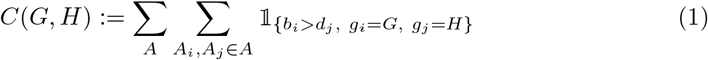

As an alternative, we used the sequence mining algorithm cSPADE [67] to determine the most frequent infection patterns from infection histories in the network. This algorithm has originally been designed to mine and classify the most frequent patterns in sets of sequences, e.g. products purchased by customers over multiple time points. In the machine learning literature, the frequency at which an item (e.g. a product, or a set of products) is encountered across a set of time-stamped customer data is referred to as ‘support’. In order to determine the most frequent multiple infection patterns at various snapshots during an epidemic, we interpreted multiple infection data as such sequences. More precisely, we sampled five random observation times *τ*_1_,… , *τ*_5_ during a simulation, and, for each host (i.e. ‘customer’), we defined the sequence of infections (i.e. ‘purchases’) by the set of genotypes the host was infected with at each observation time point. We deliberately assumed for this approach that multiple infection sequence motifs were independent snapshots of the infection state, such that re-infections and persistent infections were indistinguishable. Since our simulator allowed for multiple infections, elements of a sequence comprised the empty set, singleton sets (with only one genotype), or sets of several genotypes. The length of a motif is equivalent to the number of observations (e.g. the motif of length two < {*g*_1_, *g*_3_}, {*g*_3_} > indicated that a double infection was observed within hosts at the first time point and that a single infection was observed at the second time point). We used the R package implementation arules of the cSPADE algorithm and calculated the frequency (i.e. the ‘support’) averaged across simulation replicates and sequence motifs with minimum support of 0.02 and maximal length of 5.

## Results

### Network topology, genotype diversity, and MOI

To better understand how the topology of a contact network impacts multiple infection dynamics, we simulated epidemics by introducing simultaneously infections on random clustered graphs with distinct summary statistics. By default, we simulated multiple infections on random clustered graphs with high clustering coefficient, which we refer to as ‘clustered’ networks (see Table 1). For comparison, we considered ‘dispersed’ random clustered graphs, emulating contact networks with relatively low but variable number of contacts spread out evenly in the network due to highly over-dispersed degree and low clustering coefficient (see Table 1).

**Table 1.** Summary statistics of the two types of networks simulated. Quantities are averaged across 250 nodes and 50 stochastic replicates.

| Network type | degree | degree assortativity | degree dispersion | clustering coefficient |
| --- | --- | --- | --- | --- |
| dispersed | 4.6 | 0.58 | 51.51 | 0.13 |
| clustered | 4.1 | 0.53 | 2.36 | 0.34 |

Infection multiplicity (MOI) was highest for both homologous immunity settings, which also maintained a high diversity over the course of the simulated epidemic. In general, the network topology only had a marginal impact on MOI and diversity, but dispersed networks with high degree and low clustering coefficients maintained a higher diversity (e.g. for homologous decreasing and asymmetric settings). As expected, full cross-immunity prevented multiple infections and sharply decreased diversity, especially for clustered networks. Multiple infections could be sustained in the partial cross-immunity setting, under which the fraction of infected nodes showed a SIS-like shape due to reinfections by a subset of the circulating genotypes.

### Infection barcodes and network structure

To infer properties of the host contact network from individual infection histories, we summarised each of these through an infection barcode, i.e. a set containing all intervals of infection episodes with different parasite genotypes. In order to measure similarity (or distance) between infection barcodes of any two hosts, we used the length and frequency of overlaps between infection episodes. As detailed in the Methods, we distinguished overlaps that were genotype-specific (i.e. only for matching genotypes between two hosts) from those that were genotype-agnostic (i.e. regardless of the genotype). The resulting barcode matrices quantified the similarity (or distance) between any two hosts. These were tested for correlation to more classical matrices that capture global (shortest path between nodes) or local properties (binary adjacency) of contact networks restricted to infected individuals.

Simulation results with four different parasite genotypes show that, if the network of infected individuals was fully observed, the barcode similarity matrix significantly correlated with neighbourhood properties of the contact network given by its adjacency matrix (Figure 2). In more realistic situations, however, infected nodes would only be partially observed. Our results showed that spatial correlations were less significant as the sampling rate of network nodes decreased but remained significant for sampling proportions of 25% or more.

**Fig 1.**
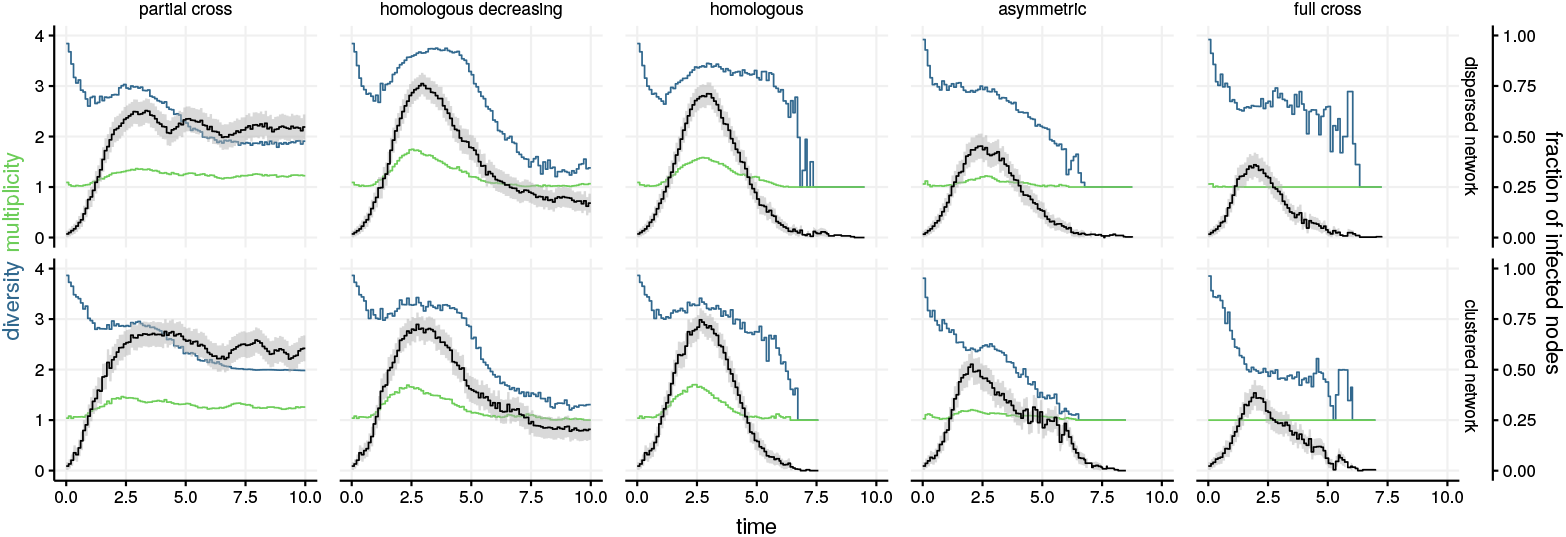
Genotype diversity index, multiplicity of infection (MOI), and epidemic prevalence as a function of network type and cross-immunity patterns. The figure shows the output of 50 stochastic epidemics with four genotypes on dispersed and clustered networks (see Table 1), and five immunological interference settings. Black lines show the time-averaged fraction of nodes infected with at least one genotype (95% confidence intervals shaded grey), blue lines show the average genotype diversity index at the population level, and green lines show the average MOI of individual nodes.

**Fig 2.**
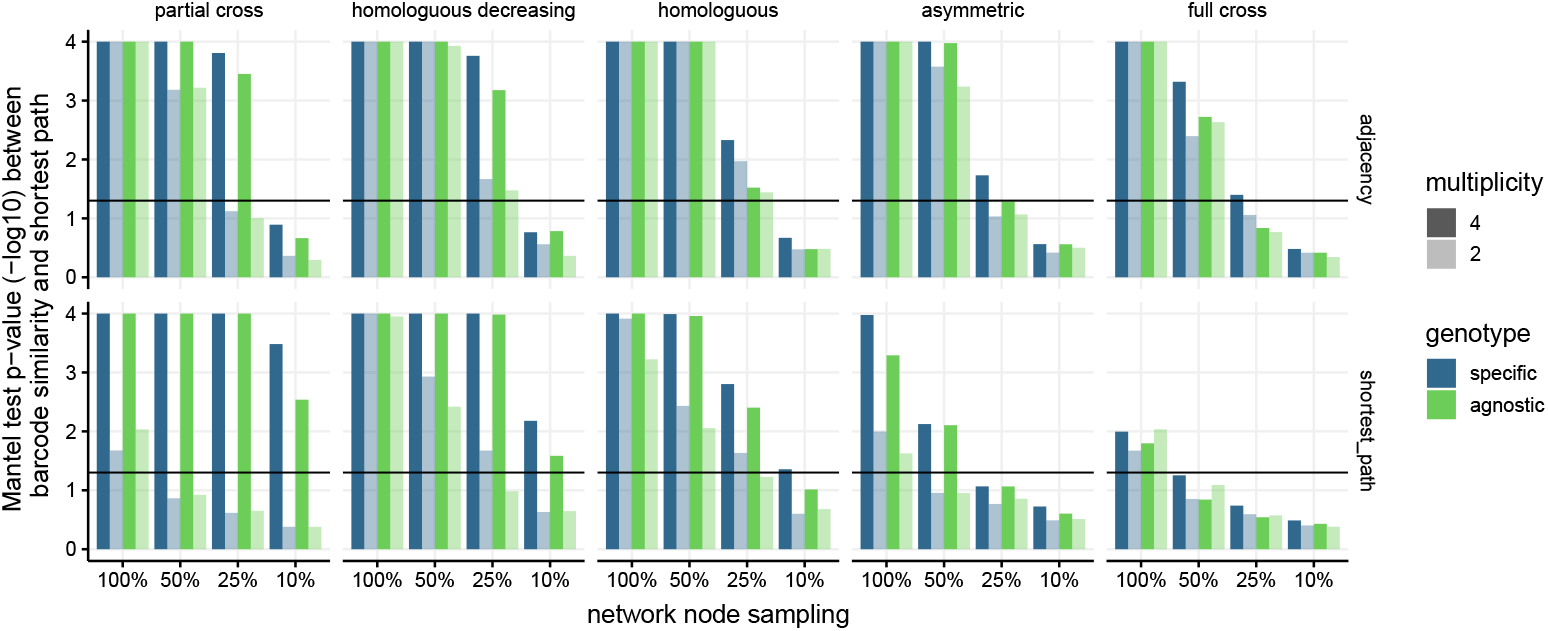
Correlations between a matrix based on infection barcodes and the network’s adjacency (top) or shortest path (bottom) matrices. For simulated epidemics on clustered networks with 2 or 4 circulating genotypes and a variety of cross-immunity settings, we tested for correlations using p-values of two-sided Mantel tests with 10^4^ permutations. For each setting, we re-sampled 20 times randomly 100, 50, 25, or 10% of the infected nodes and report the average p-value.

When pairwise comparisons between hosts were only based on the number of genotypes (green bars) but not on their nature (blue bars), our ability to detect significant correlations decreased. Unsurprisingly given the importance of multiple infection histories for our approach, decreasing the number of circulating genotypes from four (dark bars) to two (light bars) also decreased correlation significance. Cross-immunity assumptions allowing for multiple infections with highly diverse infection barcodes (e.g. full cross-immunity, asymmetric decreasing) resulted in higher significance levels. Significance levels and correlation values appeared to be higher when tested for global network properties given by the shortest path matrix (bottom), which is consistent with the barcode similarity being a quantitative measure.

We found that the correlations were highly significant even at a network sampling rate of 10% (S1 Fig), but suffered similar limitations once the number of genotypes circulating in the epidemic was reduced to two. Compared to clustered networks, we observed similar trends on dispersed networks, and correlations between barcode similarity and shortest paths tended to be higher (S2 Fig).

Overall, we found that the genotype-specific similarity index for infection barcodes was more informative than the distance metric (S3 Fig). Correlation coefficients obtained from the Mantel test were as high as 0.5 for comparisons of barcode similarity matrices with shortest path matrices regardless of network sampling rates and number of genotypes (S1 Fig).

The spatial correlations between barcode similarity indices and both local and global graph properties also translated to the individual level. From the barcode similarity matrix, we calculated each host’s connectivity (i.e. degree) in terms of infection history by summing the host’s barcode similarity with respect to all other infected hosts and normalising appropriately (see Methods). We hypothesised that the infection barcode connectivity within the network of infected hosts could inform the degree of connectivity in the contact network. First, the significant relationship between the barcode and shortest path connectivity indicated that a host with an infection history similar to those of many other hosts had also lower shortest paths to other hosts, and was hence more centrally located in the network (Fig 3).

**Fig 3.**
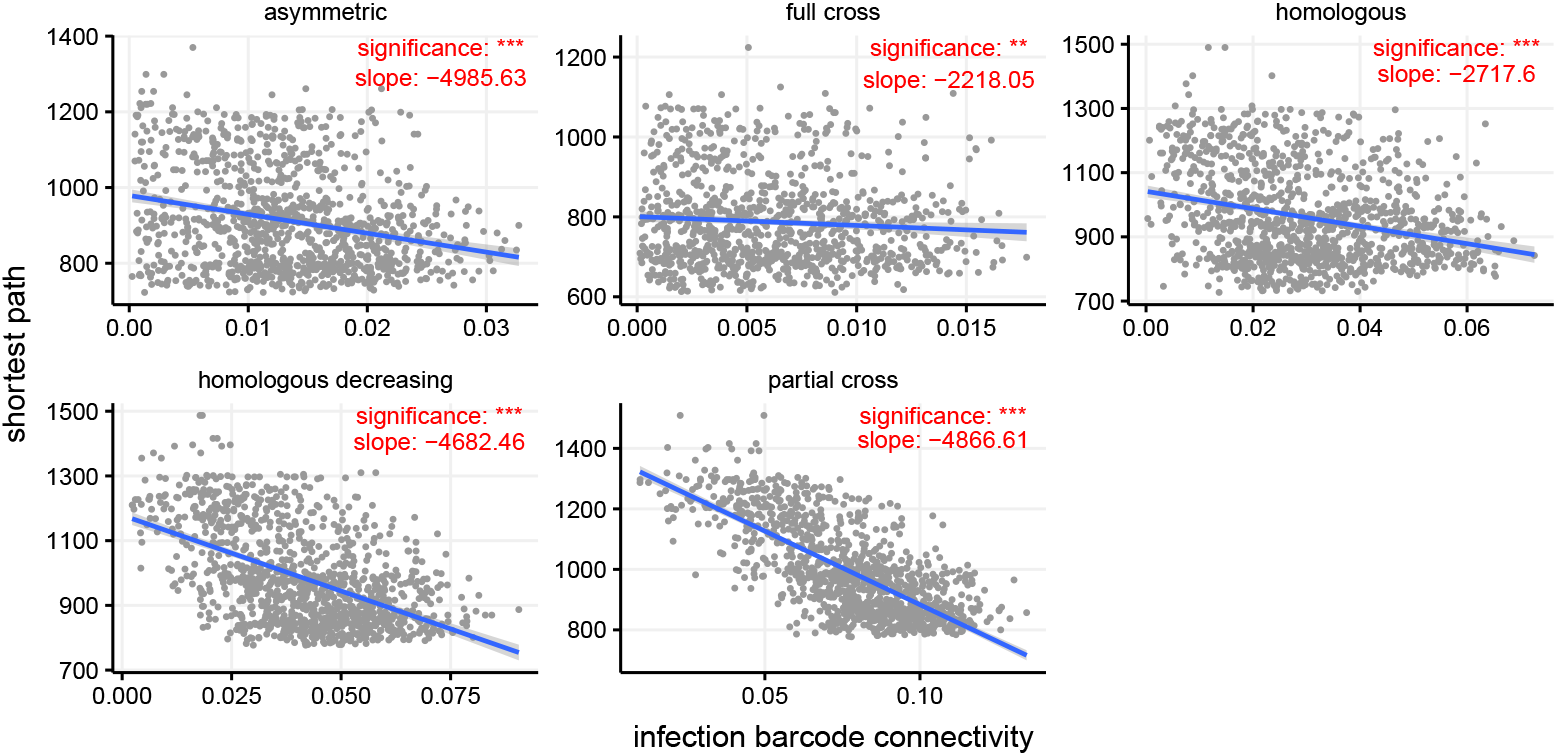
Link between the host connectivity estimated via the barcodes and that estimated via the shortest path. Signifiancy levels corresponds to p-values of a linear regression * * * : *p <* 0.001 and ** : *p <* 0.01

Multinomial regression tests showed that the odds of having higher degree in the contact network (i.e. more neighbours) increased with infection barcode connectivity. The power of the model to predict node degree was dependent on the immunity settings and highest for partial cross-immunity (Fig 4), mirroring what was seen for the continuous measure of shortest path lengths (Fig 3). For this particular immunity setting, the area under the curve was 0.74 compared to 0.6 for a multinomial model without using infection barcode connectivity indicating that predictive power increased when using multiple infection data. Similar trends, but with less predictive power, were observed for the remaining immunity settings.

**Fig 4.**
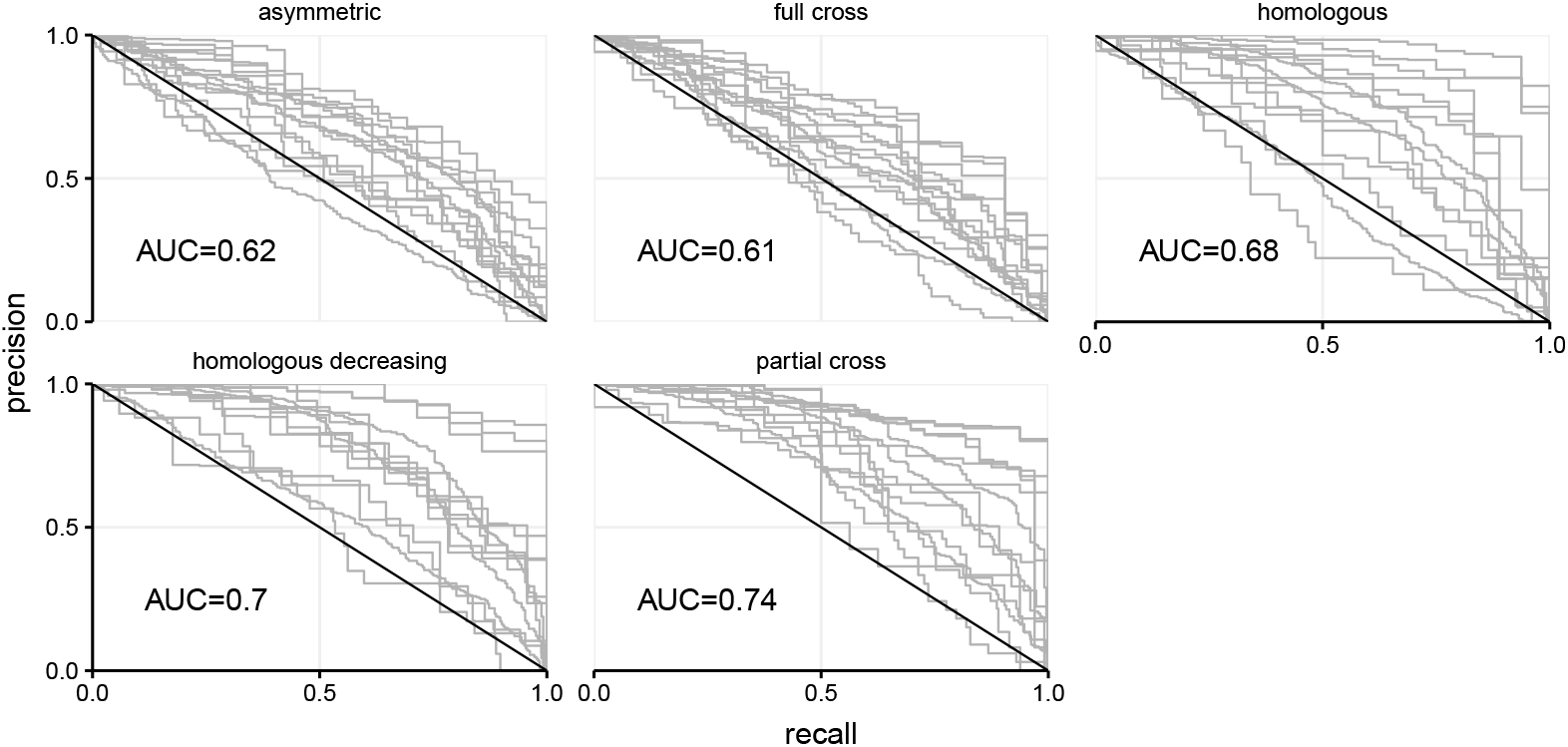
Precision-recall curve for multinomial models under five distinct immunity assumptions. The area under the curve (AUC) indicates increased (compared to random AUC of 0.5) power to classify a host’s node degree using infection barcode connectivity information.

### Infection history and immunological interference

To test whether immunological interference between genotypes could be inferred from individual infection histories, we simulated epidemics on random clustered networks with four distinct genotypes under a variety of immunity assumptions (see Methods). The co-occurrence score *C* between genotypes resembled the immunity input matrix from the simulations, in two ways. First, whenever immunity was sterile, we did not observe co-occurring genotypes (red squares in Fig 5). Second, when the probability to develop protective immunity following infection decreased, the co-occurrence score increased.

**Fig 5.**
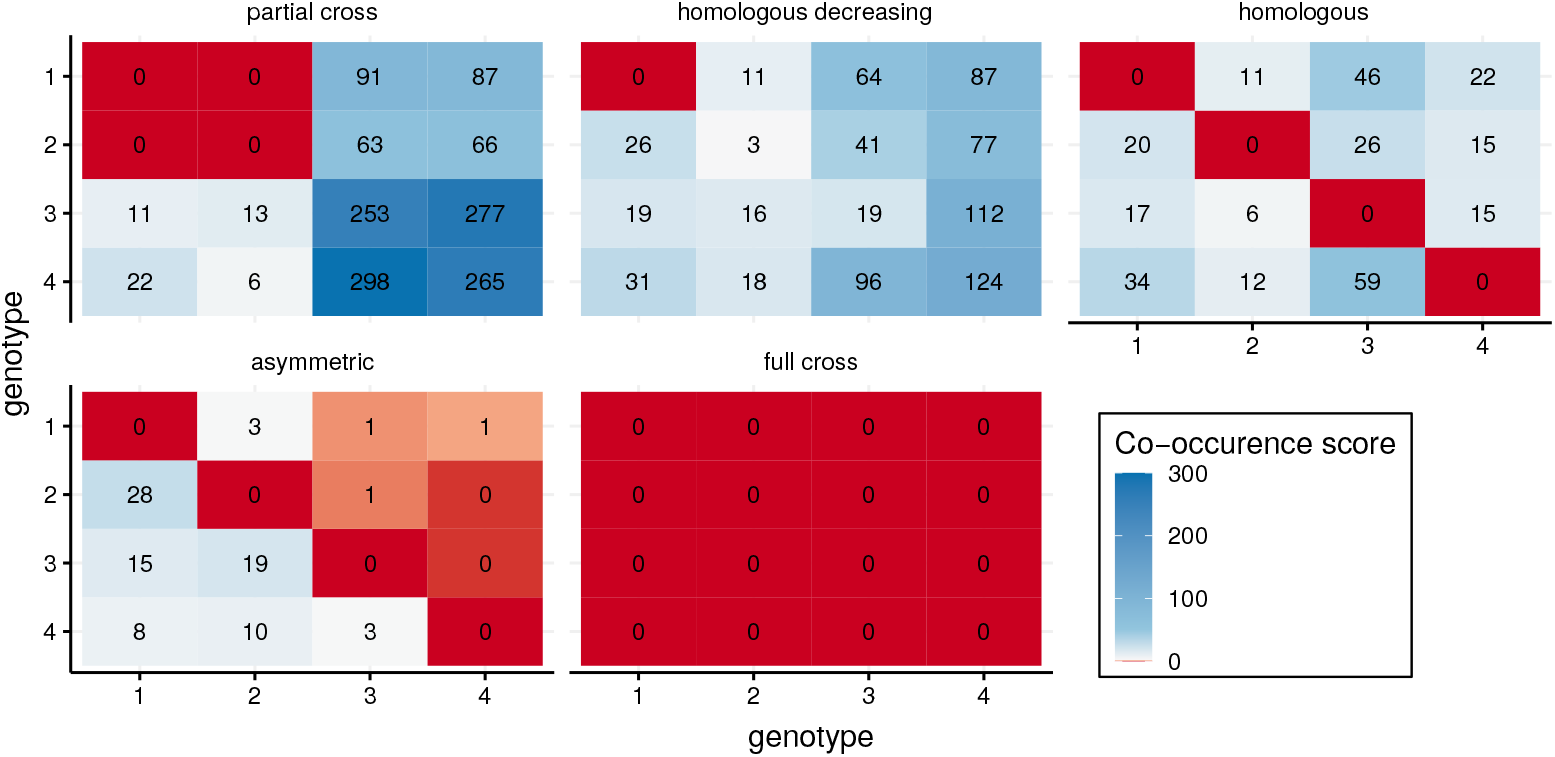
Co-occurrence score from infection barcode outputs of simulations with various assumptions on immunological interference.

The partial cross-immunity setting allowed for co-occurrence between genotypes *g*_3_ and *g*_4_, while cross-immunity between *g*_1_ and *g*_2_ lowered also possible co-occurrence with genotypes *g*_3_ and *g*_4_. Similarly, decreasing homologous immunity strongly limited co-occurrence of the same genotype, and interestingly also limited co-occurrence between genotypes which were a priori not assumed to interfere immunologically (e.g. *g*1 and *g*_4_) due to population dynamics effects. The co-occurrence score was also able to capture asymmetric immunity, stressing the possibility to estimate the order of acquisition from infection barcodes.

The sequence motif mining approach yielded infection patterns that were consistent with the immunity input (Fig 6).

**Fig 6.**
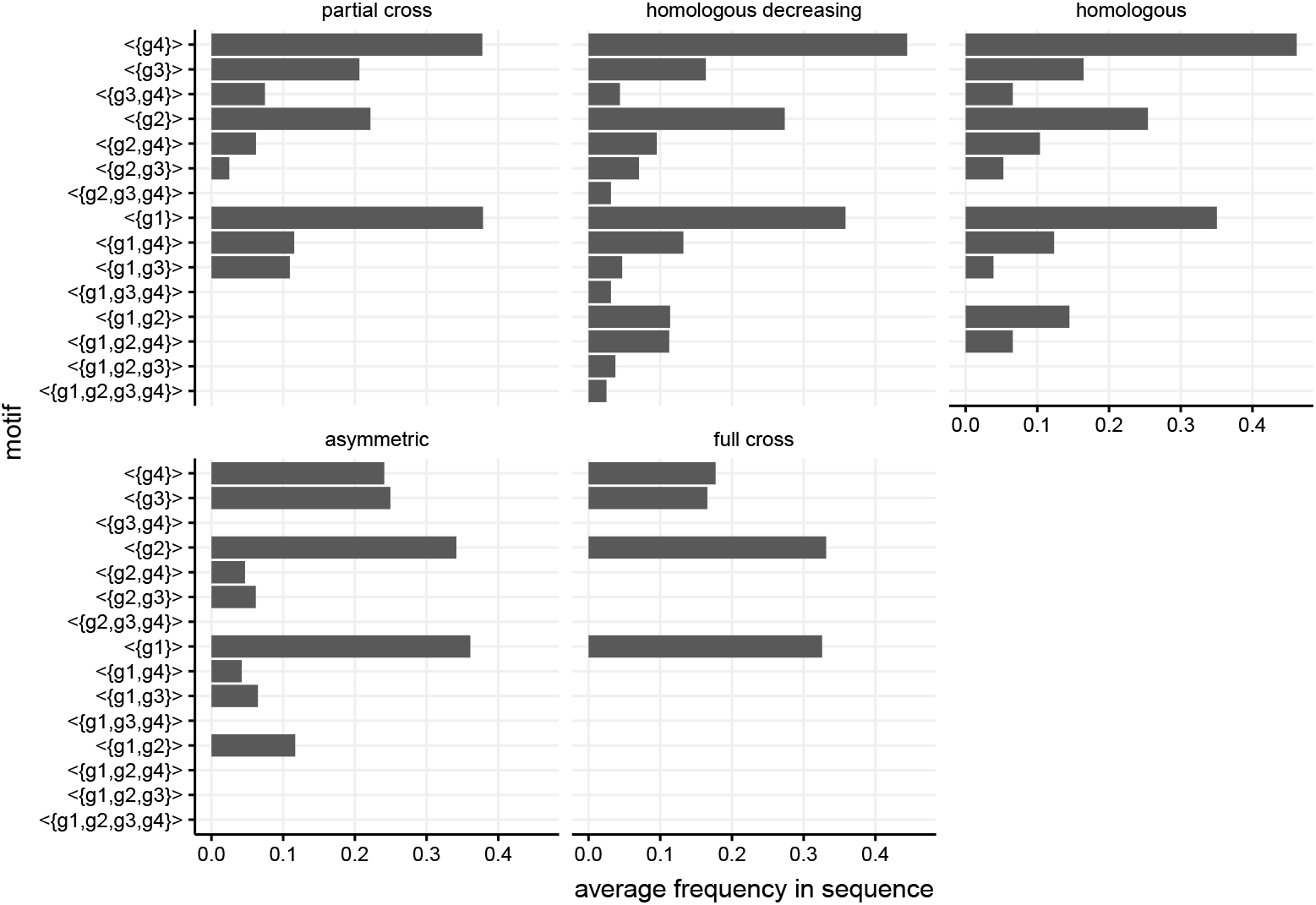
Average frequency of sequence motifs with length one, representing cross-sectional multiple infection snapshots. Samplings are performed at five random time points during the simulation.

For partial cross-immunity, multiple infections including genotypes *g*_1_ and *g*_2_ were absent from the motifs, whereas infections including the remaining genotypes were abundant with high frequency. The homologous decreasing setting was mirrored by a continuous increase in motifs from genotypes *g*_1_ to *g*_4_. In the homologous setting, motifs were dominated by single infections, whereas double infections were equally frequent between genotypes. Unsurprisingly, sterilising cross-immunity excluded all multiple infections. In order to distinguish the order of genotype acquisition, we had to consider motifs of length two (S4 Fig). In this case, differences in motif frequency with asymmetric immunity assumptions (e.g. the motif < {*g*_4_}, {*g*_1_} > was more frequent than < {*g*_1_}, {*g*_4_} >) could be detected. In general, the higher computational efficiency of sequence mining compared to the co-occurrence score was offset by the fact that, in the former, motifs are independent snapshots of the infection state, which makes reinfections and persistent infections indistinguishable.

## Discussion

Understanding the properties of host contact networks is key to predicting epidemic spread but raises many practical challenges [18]. Phylodynamic studies have shown that some of these properties can be inferred from microbial sequence data [22, 68]. Here, we use individual infection histories to gain insights into the contact network structure.

The first obstacle was to simulate epidemics on contact networks while allowing for coinfections, i.e. the simultaneous infection of hosts by multiple parasites [69]. To enable clearance events based on a host’s multiple infection history, we implemented a non-Markovian version of the Gillespie algorithm following recent developments in computational physics [36, 70]. As expected, network topology directly impacted multiple infection dynamics, with an increased level of clustering leading to higher parasite strain diversity.

Being able to simulate epidemics of multiple infections on networks, we sought to compare complex individual infection histories in order to reconstruct transmission networks. To address this issue, we captured these histories using barcodes and compared infection histories between hosts using tools from computational topology [43]. We could show that global properties of the network are correlated with these barcodes, i.e. similarity matrices inferred from the barcodes correlate with matrices inferred from the network adjacency matrix. Furthermore, individual-centred properties such as a host’s degree can also be inferred from infection barcodes.

Detecting within-host interactions between pathogens from population-level data is relevant both for persistent (*e.g.* HPV [71, 72]) and acute (*e.g.* influenzavirus [73, 74]) infections. In this simulation study, we focused on possible immunological interference between pathogens leading to exclusion mechanisms for multiple infections. We show that some patterns can be identified using multiple infection data.

Overall, we provide proof-of-principle that multiple infections and infection history can be used to gain insight into host contact network properties and immunological interference. This is biologically sound, given that infections by multiple parasite genotypes are extremely frequent, and realistic. Indeed, for many human infections, there are longitudinal follow-ups with a detailed record of parasite diversity, one of the clearest examples being human papillomaviruses [75]. Studies also exist that follow individuals in natural populations [76, 77].

Individual-based models are particularly amenable to determine new ways to analyse multiple infection data from community-based routine surveillance cohorts [73, 74]. Our model is tailored towards quantities that could actually be observed in the field, e.g. the average number of contacts and the clustering coefficient can be extracted from surveys [78]. How precise genotyping should be is a highly relevant question. Furthermore, the time component is essential to distinguish ongoing infection from reinfection. However, for many acute infectious diseases, this issue can be addressed by spacing sampling time points.

There are several limitations in our work with possible extensions. First, we only reported results obtained on random clustered networks. We also obtained similar results on other types of topologies (e.g. Erdős-Rényi, data not shown) but it would be valuable to know whether infection history is more or less informative depending on the type of network considered and if network comparison can be performed. We also assumed that the contact network was static, but in reality, it can be highly dynamic. Recent evidence suggests that these dynamics could be detected in sequence data [79] and it would be interesting to explore this with infection histories.

We assumed the life-cycle and transmission mechanisms of the parasites to be equivalent. Simulating parasite spread with distinct infectivity and clearance parameters could show the robustness of barcode metrics. Also, within-host dynamics such as viral load and immune memory could further identify relevant mechanisms for population-level dynamics of multiple infections. Following many existing studies, e.g. in phylodynamics, we used a neutrality assumption. However, genotypes are known to interact [80] and multiple infections can be a means to detect these interactions [71, 81]. In general, exploring richer life-cycles is a promising path for future studies.

We considered that infection history was based on parasite genotype presence in the host (e.g. PCR detection test) but much more data could be obtained using serologies, i.e. evidence of past presence in the host via antibody detection. An advantage of this approach is that it does not require a detailed longitudinal follow-up. However, the downside is that we then ignore the origin of the infection. Simulation studies could be instrumental in assessing our ability to infer network properties from serological data.

A separate line of future research has to do with the inference *per se*. One possibility could be to use Approximate Bayesian Computation to get a more precise idea of the accuracy of the prediction we can make on key network topology parameters. This could also allow us to compare between different classes of networks, e.g. using random forest algorithms [82].

Finally, with the increased power and decreasing cost of sequencing, it may be possible in the future to have both the information about infection history and the sequence data for multiple pathogens. It would then be worth determining whether we can get the best out of the two types of data, infection history being less precise but also less noisy than phylodynamic inference.

## Acknowledgments

This work has been funded by the European Research Council (ERC) under the European Union’s Horizon 2020 research and innovation program (EVOLPROOF, grant agreement No 648963 to SA). The authors acknowledge additional support from the CNRS, the IRD and the itrop HPC (South Green Platform in Montpellier for providing HPC resources that have contributed to the research results reported within this study (https://bioinfo.ird.fr/).

---

**Fig S1.**
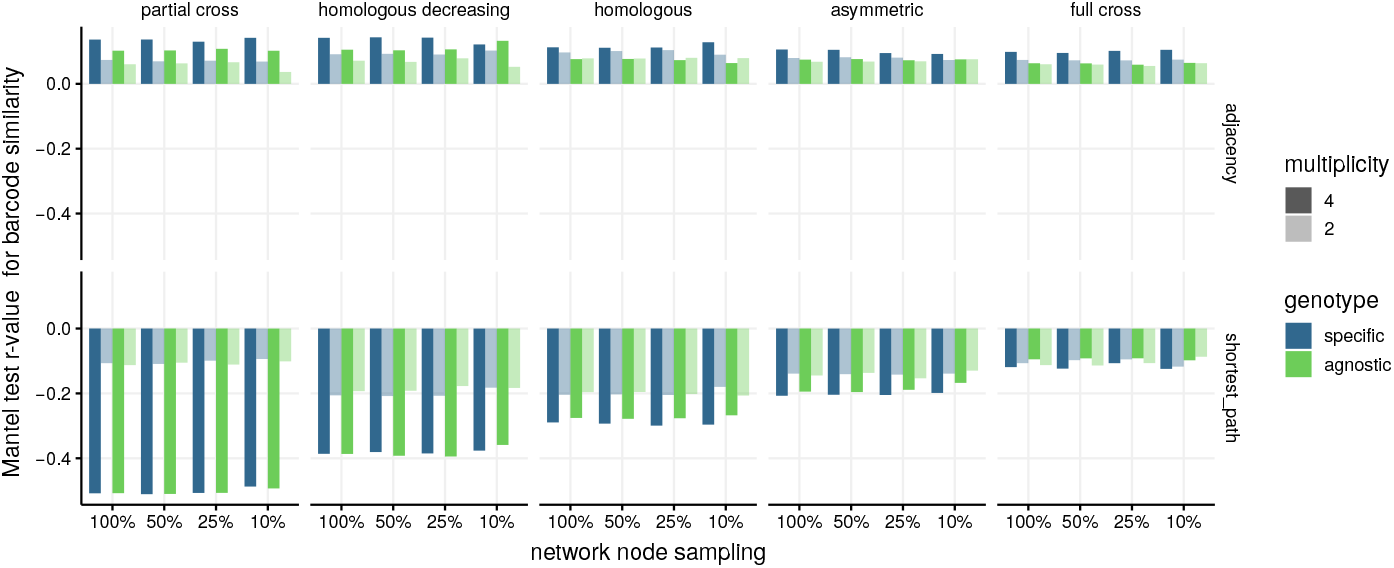
Spatial correlations. Spatial correlations between an infection barcode similarity and the network’s shortest path and adjacency matrix. We simulated epidemics with 2 or 4 circulating genotypes and for each combination of immunity setting for clustered networks, we resampled 20 times randomly 100, 50, 25 or 10% of infected nodes and report the average r-value.

**Fig S2.**
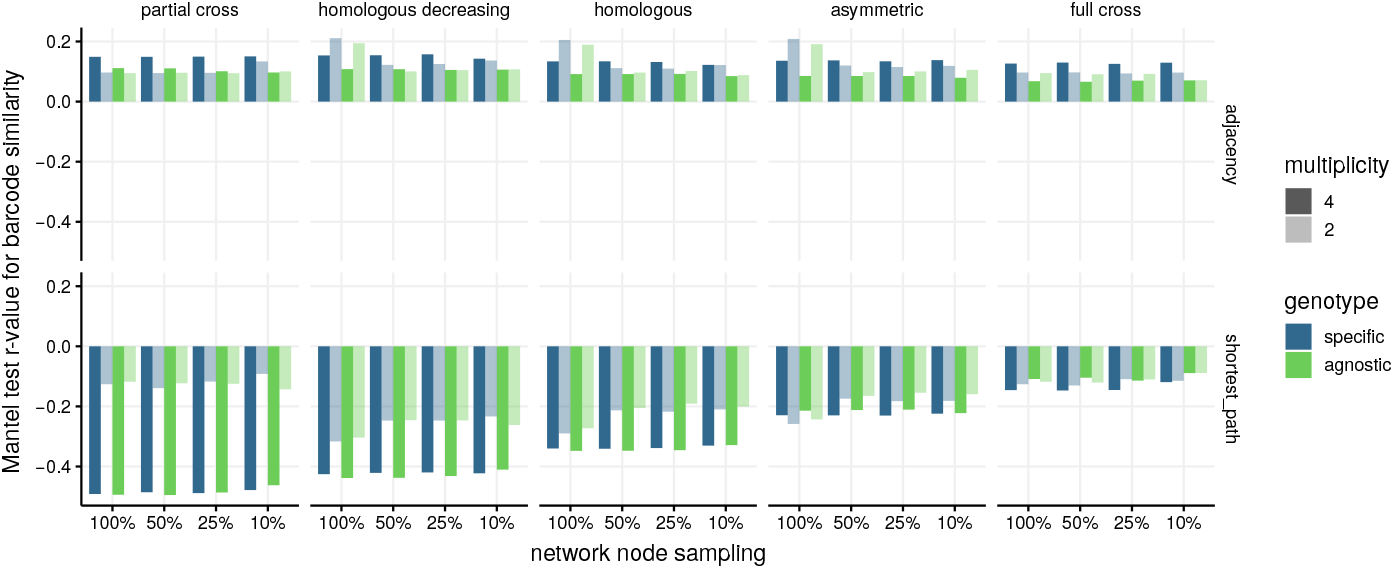
Spatial correlations for dispersed networks. Spatial correlations between an infection barcode similarity and the network’s shortest path matrix on dispersed networks. We simulated epidemics with 2 or 4 circulating genotypes and for each combination of immunity setting, we resampled 20 times randomly 100, 50, 25 or 10% of infected nodes and report the average r-value.

**Fig S3.** Spatial correlations with barcode distance. Testing for spatial correlations between an infection barcode distance and the network’s shortest path matrix on networks with average clustering coefficient of 0.34. We simulated epidemics with 2 or 4 circulating genotypes and for each combination of immunity setting and network clustering, we resampled 20 times randomly 100, 50, 25 or 10% of infected nodes and report the average p-value.

**Fig S4.** Mining for motifs of length two. Frequency of motifs with length two are displayed depending for five immunity settings, averaged across stochastic replicates.

**Fig S5.**
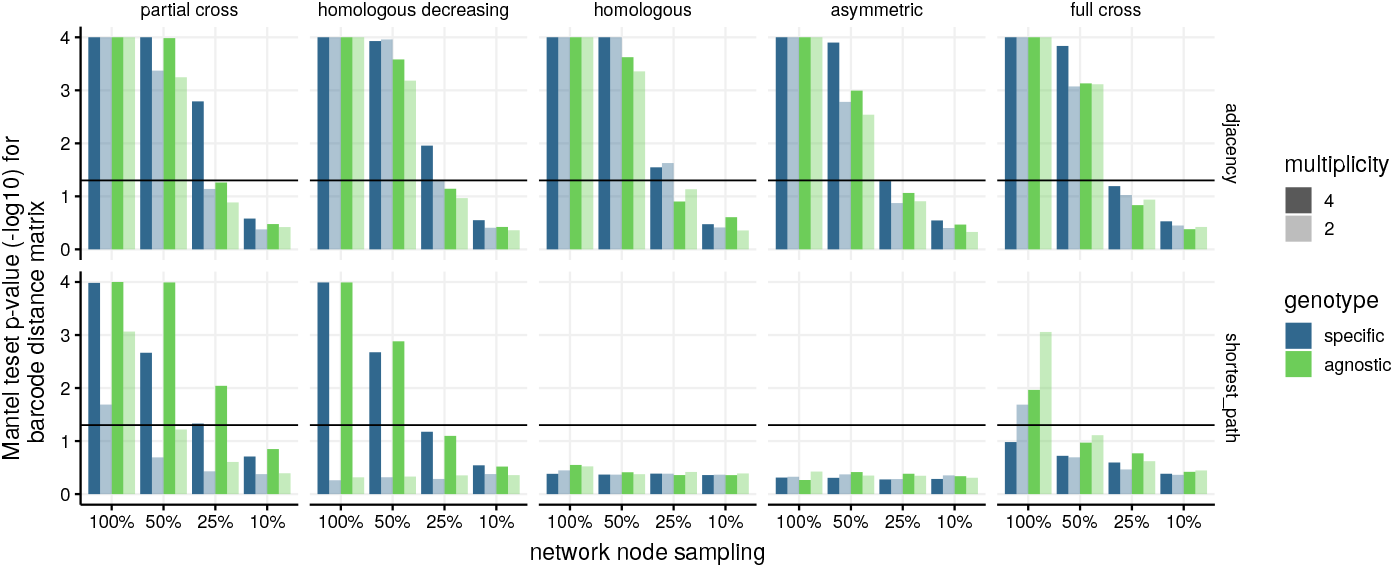
Spatial auto-correlations. Fraction of simulations with significant spatial auto-correlations in adjacency and shortest path matrices.

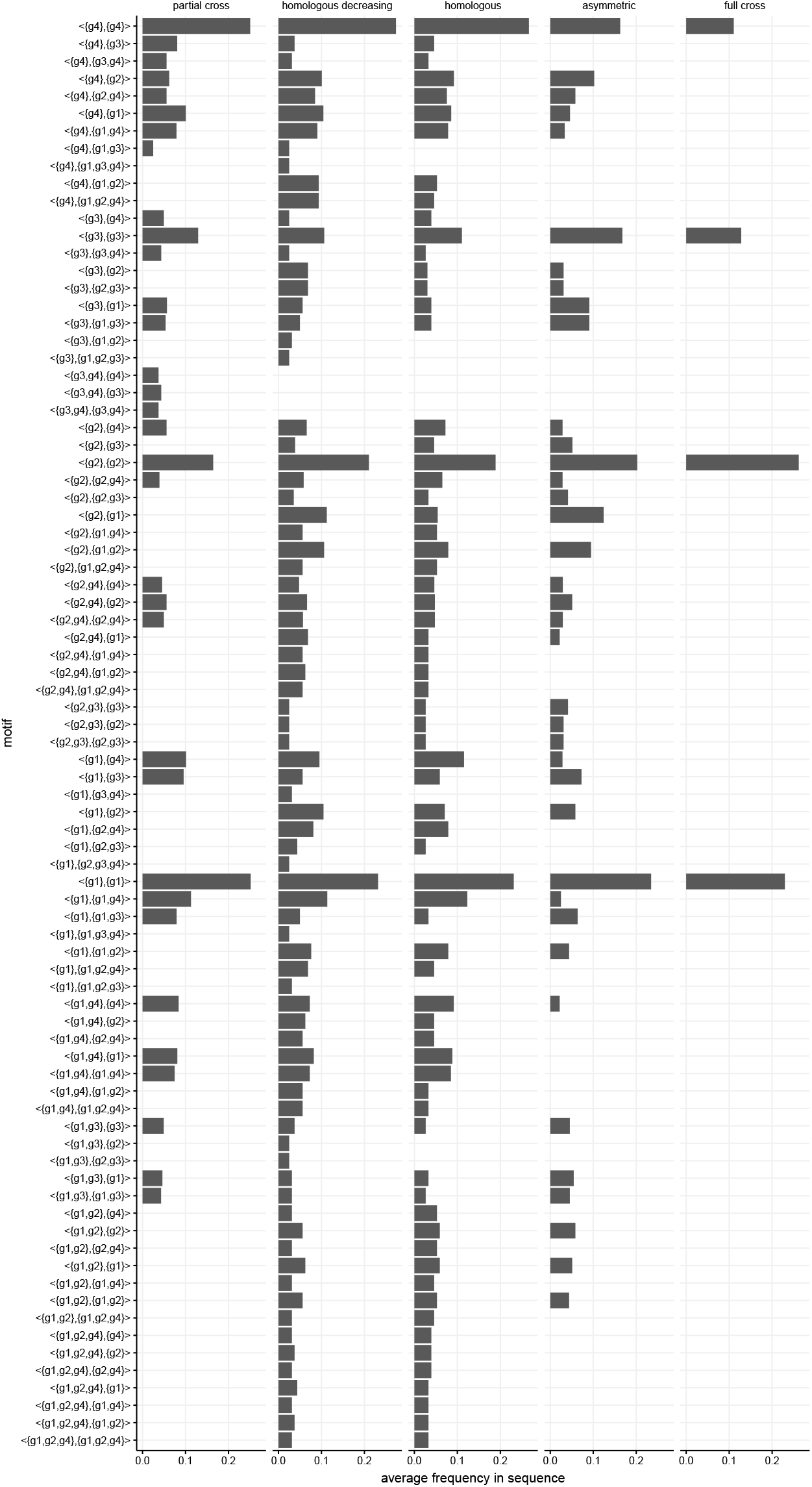

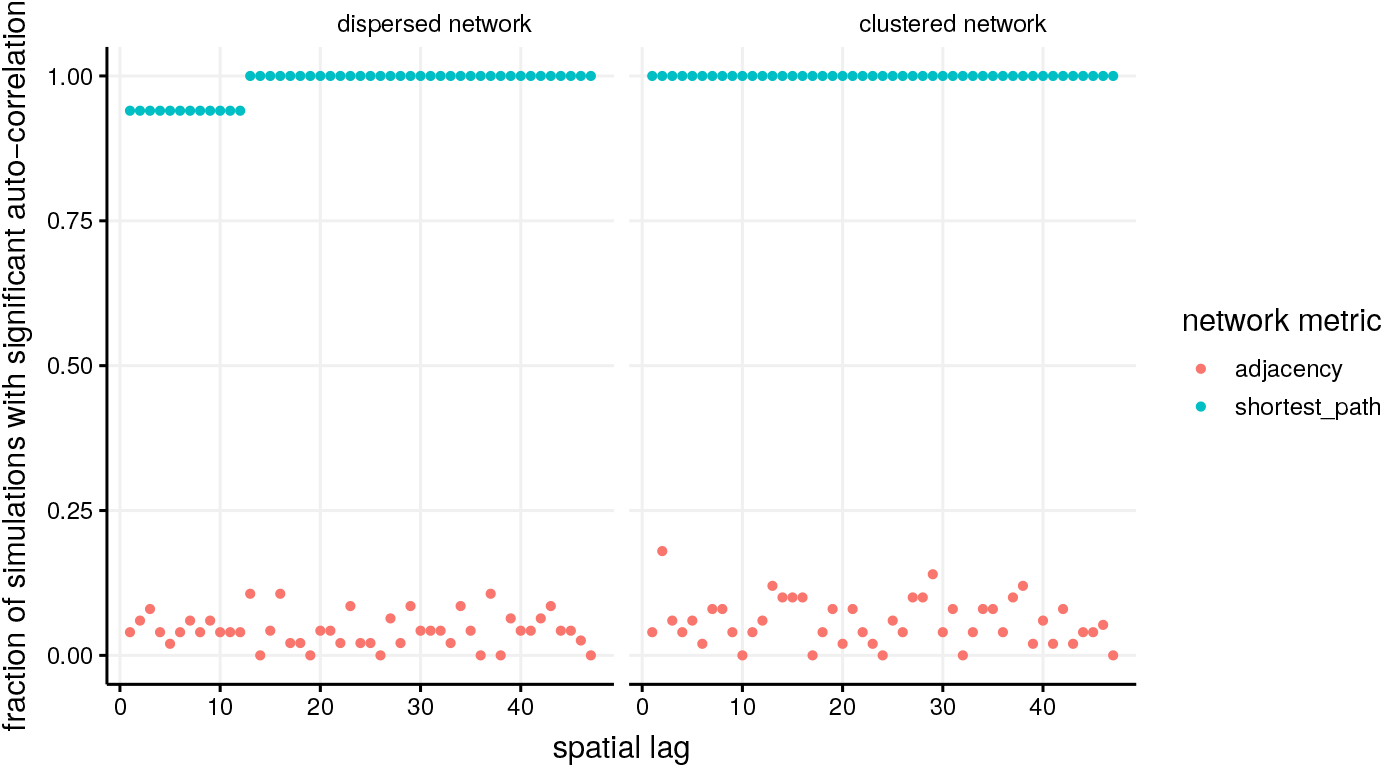

## References

1. Anderson RM, May RM. Infectious Diseases of Humans. Dynamics and Control. Oxford: Oxford University Press; 1991.

2. Diekmann O, Heesterbeek J. Mathematical epidemiology of infectious diseases: model building, analysis, and interpretation. New York: Wiley; 2000.

3. Keeling MJ, Rohani P. Modeling infectious diseases in humans and animals. Princeton University Press; 2008.

4. May RM, Anderson RM. Transmission dynamics of HIV infection. Nature. 1987;326(6109):137–42.

5. Alexander HK, Day T. Risk factors for the evolutionary emergence of pathogens. J R Soc Interface. 2010;7(51):1455–74. doi:10.1098/rsif.2010.0123.

6. Leventhal GE, Hill AL, Nowak MA, Bonhoeffer S. Evolution and emergence of infectious diseases in theoretical and real-world networks. Nat Commun. 2015;6:6101. doi:10.1038/ncomms7101.

7. Keeling MJ. The effects of local spatial structure on epidemiological invasions. Proc R Soc Lond B. 1999;266:859–867.

8. Newman MEJ. Spread of epidemic disease on networks. Phys Rev E. 2002;66(1):16128. doi:10.1103/PhysRevE.66.016128.

9. Keeling MJ, Eames KTD. Networks and epidemic models. J R Soc Interface. 2005;2(4):295–307. doi:10.1098/rsif.2005.0051.

10. Ames GM, George DB, Hampson CP, Kanarek AR, McBee CD, Lockwood DR, et al. Using network properties to predict disease dynamics on human contact networks. Proc R Soc B. 2011;278(1724):3544–3550. doi:10.1098/rspb.2011.0290.

11. Pellis L, Ball F, Bansal S, Eames K, House T, Isham V, et al. Eight challenges for network epidemic models. Epidemics. 2015;10:58–62. doi:10.1016/j.epidem.2014.07.003.

12. Hébert-Dufresne L, Althouse BM. Complex dynamics of synergistic coinfections on realistically clustered networks. Proc Natl Acad Sci USA. 2015;112(33):10551–10556. doi:10.1073/pnas.1507820112.

13. Lieberthal B, Gardner AM. Connectivity, reproduction number, and mobility interact to determine communities’ epidemiological superspreader potential in a metapopulation network. PLOS Computational Biology. 2021;17(3):e1008674. doi:10.1371/journal.pcbi.1008674.

14. Timme M, Casadiego J. Revealing networks from dynamics: an introduction. Journal of Physics A: Mathematical and Theoretical. 2014;47(34):343001. doi:10.1088/1751-8113/47/34/343001.

15. Milling C, Caramanis C, Mannor S, Shakkottai S. On identifying the causative network of an epidemic. In: 2012 50th Annual Allerton Conference on Communication, Control, and Computing (Allerton); 2012. p. 909–914.

16. Braunstein A, Ingrosso A, Muntoni AP. Network reconstruction from infection cascades. J R Soc Interface. 2019;16(151):20180844.

17. Wan X, Liu J, Cheung WK, Tong T. Inferring Epidemic Network Topology from Surveillance Data. PLoS ONE. 2014;9(6):e100661. doi:10.1371/journal.pone.0100661.

18. Eames K, Bansal S, Frost S, Riley S. Six challenges in measuring contact networks for use in modelling. Epidemics. 2015;10:72–77. doi:10.1016/j.epidem.2014.08.006.

19. Kiss IZ, Green DM, Kao RR. Disease contact tracing in random and clustered networks. Proceedings of the Royal Society B: Biological Sciences. 2005;272(1570):1407–1414. doi:10.1098/rspb.2005.3092.

20. Colizza V, Barrat A, Barthélemy M, Vespignani A. The role of the airline transportation network in the prediction and predictability of global epidemics. Proc Natl Acad Sci USA. 2006;103(7):2015–20. doi:10.1073/pnas.0510525103.

21. Eagle N, Pentland AS, Lazer D. Inferring friendship network structure by using mobile phone data. Proc Natl Acad Sci USA. 2009;106(36):15274–15278. doi:10.1073/pnas.0900282106.

22. Leventhal GE, Kouyos R, Stadler T, Wyl Vv, Yerly S, Böni J, et al. Inferring epidemic contact structure from phylogenetic trees. PLoS Comput Biol. 2012;8(3):e1002413. doi:10.1371/journal.pcbi.1002413.

23. Valdano E, Poletto C, Giovannini A, Palma D, Savini L, Colizza V. Predicting Epidemic Risk from Past Temporal Contact Data. PLoS Comput Biol. 2015;11(3):e1004152. doi:10.1371/journal.pcbi.1004152.

24. Twisk JW. Applied longitudinal data analysis for epidemiology: a practical guide. Cambridge University Press; 2013.

25. Juliano JJ, Porter K, Mwapasa V, Sem R, Rogers WO, Ariey F, et al. Exposing malaria in-host diversity and estimating population diversity by capture-recapture using massively parallel pyrosequencing. Proc Natl Acad Sci USA. 2010;107(46):20138–43. doi:10.1073/pnas.1007068107.

26. Chaturvedi AK, Katki HA, Hildesheim A, Rodríguez AC, Quint W, Schiffman M, et al. Human Papillomavirus Infection with Multiple Types: Pattern of Coinfection and Risk of Cervical Disease. J Infect Dis. 2011;203:910–920. doi:10.1093/infdis/jiq139.

27. Shaw DJ, Dobson AP. Patterns of macroparasite abundance and aggregation in wildlife populations: a quantitative review. Parasitology. 1995;111(S1):S111–S133. doi:10.1017/S0031182000075855.

28. Alizon S, Murall CL, Saulnier E, Sofonea MT. Detecting within-host interactions from genotype combination prevalence data. Epidemics. 2019;29:100349. doi:10.1016/j.epidem.2019.100349.

29. Opatowski L, Baguelin M, Eggo RM. Influenza interaction with cocirculating pathogens and its impact on surveillance, pathogenesis, and epidemic profile: A key role for mathematical modelling. PLOS Pathogens. 2018;14(2):e1006770. doi:10.1371/journal.ppat.1006770.

30. Buckee CO, Koelle K, Mustard MJ, Gupta S. The effects of host contact network structure on pathogen diversity and strain structure. Proceedings of the National Academy of Sciences. 2004;101(29):10839–10844. doi:10.1073/pnas.0402000101.

31. Miller JC. Cocirculation of infectious diseases on networks. Phys Rev E Stat Nonlin Soft Matter Phys. 2013;87(6):060801.

32. Poletto C, Meloni S, Colizza V, Moreno Y, Vespignani A. Host Mobility Drives Pathogen Competition in Spatially Structured Populations. PLoS Computational Biology. 2013;9(8):e1003169. doi:10.1371/journal.pcbi.1003169.

33. Poletto C, Meloni S, Metre AV, Colizza V, Moreno Y, Vespignani A. Characterising two-pathogen competition in spatially structured environments. Scientific Reports. 2015;5(1). doi:10.1038/srep07895.

34. Sahneh FD, Scoglio C. Competitive epidemic spreading over arbitrary multilayer networks. Physical Review E. 2014;89(6). doi:10.1103/physreve.89.062817.

35. Pinotti F, Fleury É, Guillemot D, Böelle PY, Poletto C. Host contact dynamics shapes richness and dominance of pathogen strains. 2018;doi:10.1101/428185.

36. Boguña M, Lafuerza LF, Toral R, Serrano MA. Simulating non-Markovian stochastic processes. Phys Rev E. 2014;90(4):042108. doi:10.1103/PhysRevE.90.042108.

37. Bailey NTJ. ON ESTIMATING THE LATENT AND INFECTIOUS PERIODS OF MEASLES. Biometrika. 1956;43(1-2):15–22. doi:10.1093/biomet/43.1-2.15.

38. Eichner M, Dietz K. Transmission Potential of Smallpox: Estimates Based on Detailed Data from an Outbreak. American Journal of Epidemiology. 2003;158(2):110–117. doi:10.1093/aje/kwg103.

39. Sama W, Dietz K, Smith T. Distribution of survival times of deliberate Plasmodium falciparum infections in tertiary syphilis patients. Transactions of the Royal Society of Tropical Medicine and Hygiene. 2006;100(9):811–816. doi:10.1016/j.trstmh.2005.11.001.

40. Katki HA, Cheung LC, Fetterman B, Castle PE, Sundaram R. A joint model of persistent human papilloma virus infection and cervical cancer risk: implications for cervical cancer screening. Journal of the Royal Statistical Society: Series A (Statistics in Society). 2015;178(4):903–923. doi:10.1111/rssa.12101.

41. Bretscher MT, Maire N, Chitnis N, Felger I, Owusu-Agyei S, Smith T. The distribution of Plasmodium falciparum infection durations. Epidemics. 2011;3(2):109–118. doi:10.1016/j.epidem.2011.03.002.

42. Cowling BJ, Jin L, Lau EH, Liao Q, Wu P, Jiang H, et al. Comparative epidemiology of human infections with avian influenza A H7N9 and H5N1 viruses in China: a population-based study of laboratory-confirmed cases. The Lancet. 2013;382(9887):129–137. doi:10.1016/s0140-6736(13)61171-x.

43. Edelsbrunner H, Harer J. Computational Topology - an Introduction. American Mathematical Society; 2010.

44. Keeling MJ. Disease Extinction and Community Size: Modeling the Persistence of Measles. Science. 1997;275(5296):65–67. doi:10.1126/science.275.5296.65.

45. Lloyd AL. Realistic Distributions of Infectious Periods in Epidemic Models: Changing Patterns of Persistence and Dynamics. Theoretical Population Biology. 2001;60(1):59–71. doi:10.1006/tpbi.2001.1525.

46. Gough KJ. The estimation of latent and infectious periods. Biometrika. 1977;64(3):559–565. doi:10.1093/biomet/64.3.559.

47. Greenhalgh S, Day T. Time-varying and state-dependent recovery rates in epidemiological models. Infectious Disease Modelling. 2017;2(4):419–430. doi:10.1016/j.idm.2017.09.002.

48. Fox A, Mai LQ, Thanh LT, Wolbers M, Hang NLK, Thai PQ, et al. Hemagglutination inhibiting antibodies and protection against seasonal and pandemic influenza infection. Journal of Infection. 2015;70(2):187–196. doi:10.1016/j.jinf.2014.09.003.

49. Bansal S, Khandelwal S, Meyers LA. Exploring biological network structure with clustered random networks. BMC Bioinformatics. 2009;10(1). doi:10.1186/1471-2105-10-405.

50. Miller JC. Percolation and epidemics in random clustered networks. Physical Review E. 2009;80(2). doi:10.1103/physreve.80.020901.

51. Newman MEJ. Random Graphs with Clustering. Physical Review Letters. 2009;103(5). doi:10.1103/physrevlett.103.058701.

52. Hagberg AA, Schult DA, Swart PJ. Exploring Network Structure, Dynamics, and Function using NetworkX. In: Varoquaux G, Vaught T, Millman J, editors. Proceedings of the 7th Python in Science Conference. Pasadena, CA USA; 2008. p. 11 – 15.

53. Newman MEJ. Assortative Mixing in Networks. Physical Review Letters. 2002;89(20). doi:10.1103/physrevlett.89.208701.

54. Gillespie DT. A general method for numerically simulating the stochastic time evolution of coupled chemical reactions. Journal of Computational Physics. 1976;22(4):403–434. doi:10.1016/0021-9991(76)90041-3.

55. Gillespie DT. Exact stochastic simulation of coupled chemical reactions. The Journal of Physical Chemistry. 1977;81(25):2340–2361. doi:10.1021/j100540a008.

56. Boguña M, Lafuerza LF, Toral R, Serrano MA. Simulating non-Markovian stochastic processes. Phys Rev E Stat Nonlin Soft Matter Phys. 2014;90(4):042108.

57. Morris EK, Caruso T, Buscot F, Fischer M, Hancock C, Maier TS, et al. Choosing and using diversity indices: insights for ecological applications from the German Biodiversity Exploratories. Ecology and Evolution. 2014;4(18):3514–3524. doi:10.1002/ece3.1155.

58. Chazal F, Michel B. An introduction to Topological Data Analysis: fundamental and practical aspects for data scientists. ArXiv e-prints. 2017;.

59. Carlsson G, Zomorodian A, Collins A, Guibas L. Persistence Barcodes for Shapes. In: Proceedings of the 2004 Eurographics/ACM SIGGRAPH Symposium on Geometry Processing. SGP ’04. New York, NY, USA:ACM; 2004. p. 124–135. Available from: http://doi.acm.org/10.1145/1057432.1057449.

60. Diniz-Filho JA, Soares TN, Lima JS, Dobrovolski R, Landeiro VL, de Campos Telles MP, et al. Mantel test in population genetics. Genet Mol Biol. 2013;36(4):475–485.

61. Goslee SC, Urban DL. The ecodist package for dissimilarity-based analysis of ecological data. Journal of Statistical Software. 2007;22:1–19.

62. Guillot G, Rousset F. Dismantling the Mantel tests. Methods in Ecology and Evolution. 2013;4(4):336–344. doi:10.1111/2041-210x.12018.

63. Legendre P, Fortin MJ, Borcard D. Should the Mantel test be used in spatial analysis? Methods in Ecology and Evolution. 2015;6(11):1239–1247. doi:10.1111/2041-210x.12425.

64. Legendre P, Fortin MJ. Comparison of the Mantel test and alternative approaches for detecting complex multivariate relationships in the spatial analysis of genetic data. Molecular Ecology Resources. 2010;10(5):831–844. doi:10.1111/j.1755-0998.2010.02866.x.

65. Fujita A, Takahashi DY, Balardin JB, Vidal MC, Sato JR. Correlation between graphs with an application to brain network analysis. Computational Statistics & Data Analysis. 2017;109:76–92. doi:10.1016/j.csda.2016.11.016.

66. Hand DJ, Till RJ. Machine Learning. 2001;45(2):171–186. doi:10.1023/a:1010920819831.

67. Zaki MJ. Machine Learning. 2001;42(1/2):31–60. doi:10.1023/a:1007652502315.

68. Rasmussen DA, Kouyos R, Günthard HF, Stadler T. Phylodynamics on local sexual contact networks. PLOS Computational Biology. 2017;13(3):e1005448. doi:10.1371/journal.pcbi.1005448.

69. Sofonea MT, Alizon S, Michalakis Y. Exposing the diversity of multiple infection patterns. Journal of Theoretical Biology. 2017;419:278–289. doi:10.1016/j.jtbi.2017.02.011.

70. Vestergaard CL, Génois M. Temporal Gillespie Algorithm: Fast Simulation of Contagion Processes on Time-Varying Networks. PLOS Computational Biology. 2015;11(10):e1004579. doi:10.1371/journal.pcbi.1004579.

71. Alizon S, Murall CL, Saulnier E, Sofonea MT. Detecting within-host interactions from genotype combination prevalence data. Epidemics. 2019;29:100349. doi:10.1016/j.epidem.2019.100349.

72. Hamelin FM, Allen LJS, Bokil VA, Gross LJ, Hilker FM, Jeger MJ, et al. Coinfections by noninteracting pathogens are not independent and require new tests of interaction. PLOS Biology. 2019;17(12):e3000551. doi:10.1371/journal.pbio.3000551.

73. Chu HY, Boeckh M, Englund JA, Famulare M, Lutz B, Nickerson DA, et al. The Seattle Flu Study: a multiarm community-based prospective study protocol for assessing influenza prevalence, transmission and genomic epidemiology. BMJ Open. 2020;10(10):e037295. doi:10.1136/bmjopen-2020-037295.

74. Hoa LNM, Sullivan SG, Mai LQ, Khvorov A, Phuong HVM, Hang NLK, et al. Influenza A(H1N1)pdm09 But Not A(H3N2) Virus Infection Induces Durable Seroprotection: Results From the Ha Nam Cohort. The Journal of Infectious Diseases. 2020;doi:10.1093/infdis/jiaa293.

75. Brotman RM, Shardell MD, Gajer P, Tracy JK, Zenilman JM, Ravel J, et al. Interplay Between the Temporal Dynamics of the Vaginal Microbiota and Human Papillomavirus Detection. Journal of Infectious Diseases. 2014;210(11):1723–1733. doi:10.1093/infdis/jiu330.

76. Hayward AD, Nussey DH, Wilson AJ, Berenos C, Pilkington JG, Watt KA, et al. Natural Selection on Individual Variation in Tolerance of Gastrointestinal Nematode Infection. PLoS Biol. 2014;12(7):e1001917. doi:10.1371/journal.pbio.1001917.

77. Rynkiewicz EC, Pedersen AB, Fenton A. An ecosystem approach to understanding and managing within-host parasite community dynamics. Trends in Parasitology. 2015;31(5):212–221. doi:10.1016/j.pt.2015.02.005.

78. Horby P, Mai LQ, Fox A, Thai PQ, Yen NTT, Thanh LT, et al. The Epidemiology of Interpandemic and Pandemic Influenza in Vietnam, 2007–2010. American Journal of Epidemiology. 2012;175(10):1062–1074. doi:10.1093/aje/kws121.

79. Metzig C, Ratmann O, Bezemer D, Colijn C. Phylogenies from dynamic networks. PLOS Computational Biology. 2019;15(2):e1006761. doi:10.1371/journal.pcbi.1006761.

80. Mideo N. Parasite adaptations to within-host competition. Trends in Parasitology. 2009;25(6):261–268. doi:10.1016/j.pt.2009.03.001.

81. Man I, Auranen K, Wallinga J, Bogaards JA. Capturing multiple-type interactions into practical predictors of type replacement following human papillomavirus vaccination. Philosophical Transactions of the Royal Society B: Biological Sciences. 2019;374(1773):20180298. doi:10.1098/rstb.2018.0298.

82. Pudlo P, Marin JM, Estoup A, Cornuet JM, Gautier M, Robert CP. Reliable ABC model choice via random forests. Bioinformatics. 2015;32(6):859–866. doi:10.1093/bioinformatics/btv684.

